# Most metagenomic vanT calls are not vancomycin resistance genes

**DOI:** 10.64898/2026.10.22.760690

**Authors:** Prateek Shetty, Gergely Maroti, Roland Wirth

## Abstract

Metagenomic resistome surveys frequently report *vanT*_*G*_ as the dominant glycopeptide resistance gene, but these calls rely on homology without verifying catalytic residues or operon context. By analyzing 107,008 metagenome-assembled genomes, we demonstrate that only 3.7% of predicted *vanT*_*G*_ genes possess the required active-site substitution, with the vast majority being misannotated alanine racemases. Incorporating structural and genomic context corrections reveals a novel split-architecture lineage in gut-associated Erysipelotrichales missed by current profiling tools.

## Main

Surveys of antimicrobial resistance genes in metagenome-assembled genomes are one method to track the spread of resistance determinants^1–4^. These rankings rest on homology-based gene calls, and the calls are rarely extended to ask whether the encoded protein carries the catalytically important residues, the domains, or the operon context that the function requires.

We used CARD’s^6^ RGI tool on 107,008 SPIRE^5^ metagenome-assembled genomes (MAGs) and identified that 77,584 MAGs carry at least one resistance gene call, with glycopeptide resistance being the most prevalent class, appearing in 65,130 MAGs (Fig. 1a). This abundance is driven by one gene: *vanT* within the *vanG* operon (*vanT*_*G*_ hereafter). *vanT*_*G*_ is called 49,389 times, and is present in 46,168 genomes; 43% of everything we screened (Fig. 1b). Any survey reporting glycopeptide resistance as the dominant class in environmental metagenomes is primarily reporting this gene. A globally distributed wastewater resistome survey covering 142 treatment plants on six continents names *vanT*_*G*_ as the third most abundant resistance gene worldwide at 11.4% of all calls, and glycopeptide as the second most abundant class at 24.5%^1^. The same pattern recurs across unrelated environments and is not specific to wastewater. In a eutrophic plateau lake, *vanT*_*G*_ was the single most abundant resistance gene in both the dry and wet seasons^11^. It ranks second, behind only *adeF*, in hospital sink-drain biofilms^12^, and is predominant in infant gut metagenomes^13^, a core transcriptionally active gene across municipal wastewater treatment^14^, and detected in every participant of a longitudinal human gut study^15^.

**Fig. 1.**
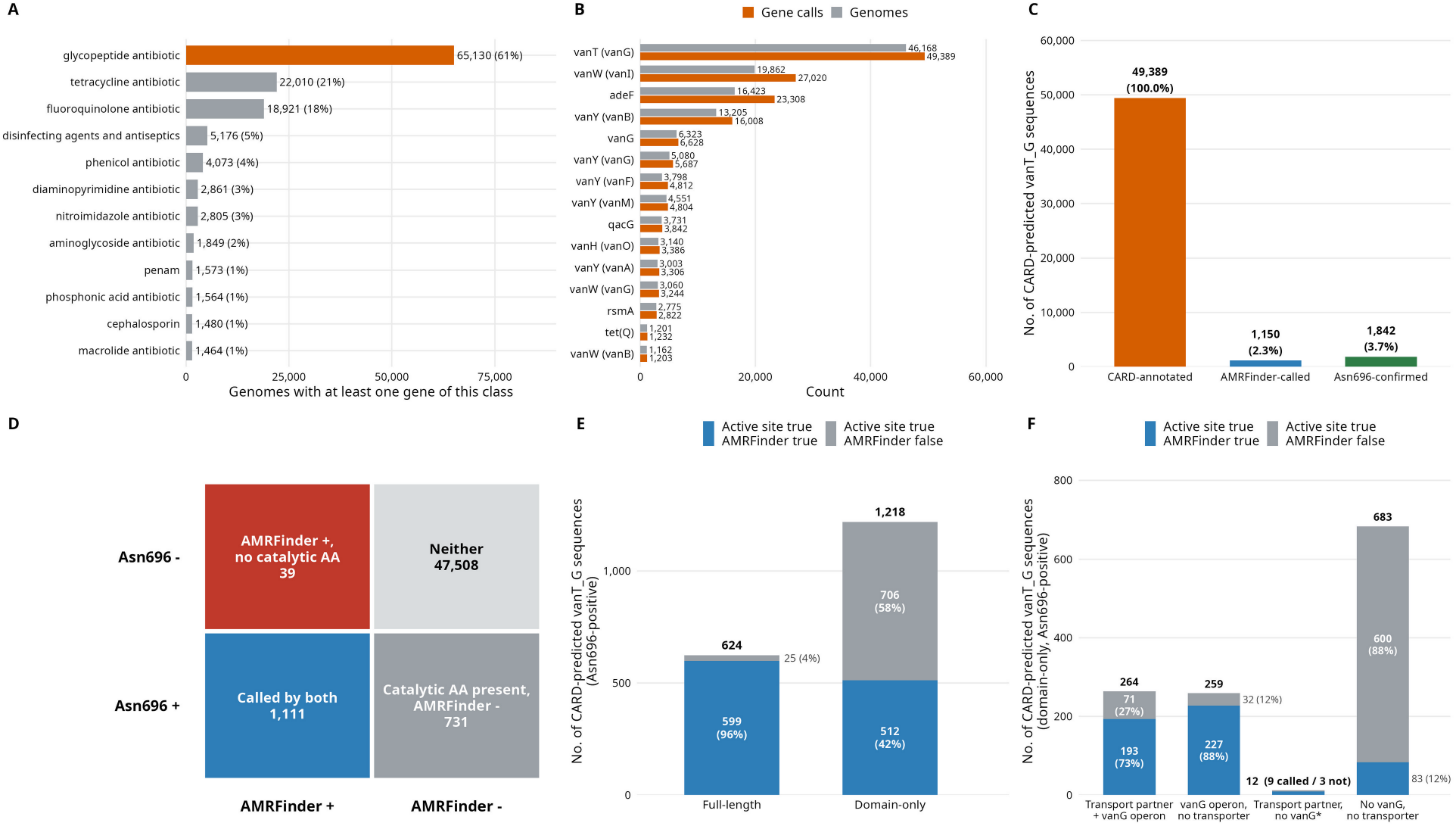
Most metagenomic *vanT*_*G*_ calls lack the catalytic residue. A) Number of MAGs carrying at least one call of each of the twelve most prevalent drug classes, across 107,008 SPIRE species-representative genomes. The number of genomes carrying glycopeptide antibiotic resistance is highlighted in orange and the rest are in grey. B) The fifteen most-called AMR genes. The number of calls is in orange and the number of genomes carrying them is in grey. C) The three values of the three prediction tools calling *vanT*_*G*_. CARD calls are shown in orange (49,389), AMRFinderPlus calls are shown in blue (1,150) and number of sequences carrying Asn696 is shown in green (1,842). D) Agreement between AMRFinderPlus and the active-site test, as the four quadrants. E) The 1,842 active-site-positive loci by domain architecture separated as full-length versus catalytic domain only. F) The 1,218 domain-only active-site-positive loci grouped by genomic context (adjacent to a gene carrying transmembrane transport domain and/or vanG ligase within 10 kb) and by AMRFinderPlus call.

Vancomycin binds the D-alanyl-D-alanine terminus of the peptidoglycan precursor, blocking the crosslinking that grows cell wall, eventually leading to cell lysis. Every characterised van operon overcomes the antibiotic in the same way, by remodelling the cell wall precursors, so the antibiotic no longer binds. The *vanA, vanB, vanD, vanF, vanI* and *vanM* operons substitute D-alanine with D-lactate, reducing vancomycin affinity to the cell wall by roughly a thousandfold and conferring high-level resistance. The other operons include *vanC, vanE, vanG* and *vanL*, which substitute D-alanine with D-serine (Supplementary Table 1). This substitution reduces the vancomycin affinity by around sixfold, and confer low-level resistance.

Supplying the D-serine is *vanT*’s role. VanT_G_ is a bifunctional membrane serine racemase with an N-terminal transmembrane transport domain and a C-terminal catalytic domain that converts L-serine to D-serine for incorporation into the modified terminus. The catalytic domain belongs to the alanine racemase family and closely resembles the housekeeping alanine racemases carried by essentially every bacterium. The alanine racemases supply D-alanine for the wild-type wall. The VanT_G_ active site contains four amino acids that form a unique hydrogen-bonded network not observed in typical alanine racemases: Tyr543, Ser567, Thr695, and Asn696. While all four residues are conserved among VanT enzymes, Asn696 is the most critical determinant in L-serine selectivity. In the VanT_G_ reference structure (PDB 4ECL), this asparagine occupies the position held by tyrosine in alanine racemases. Reverting it experimentally reduces L-serine turnover roughly thirtyfold while activity on L-alanine remains intact^9^. We refer to this residue as Asn696 throughout this work; because the equivalent position carries a different coordinate in every homologue, it is located throughout this work by alignment to the reference HMM profiles.

A gene that confers resistance must do three things: carry the catalytic residue, import the substrate, and sit beside the ligase that uses the product. Applying the active-site requirement to our CARD predicted *vanT*_*G*_ sequences reveals a sharp drop from the 49,389 CARD predictions: only 1,842 sequences (3.7%) actually carry Asn696 (Fig. 1c). The remainder are not variants of the same enzyme, as 87.8% of the 47,223 aligned CARD predicted *vanT*_*G*_ sequences carry tyrosine, the canonical residue of housekeeping alanine racemases. As an independent check, we also ran AMRFinderPlus^7,8^ on the CARD-predicted *vanT*_*G*_ calls. Because AMRFinderPlus uses manually curated hidden Markov models with fixed trusted cutoffs rather than sequence-identity thresholds, its predictions drop to a similarly small fraction of 1,150 calls (2.3%; Fig. 1c). Comparing the two approaches directly shows that they agree on 1,111 calls. Most importantly, AMRFinderPlus calls 39 sequences that lack Asn696 at the catalytic site and does not call 731 active-site-positive loci (Fig. 1d).

Among the 1,842 active-site-positive loci, only 624 are full-length genes with both domains, whereas 1,218 encode the catalytic domain alone. AMRFinderPlus calls 96% of the full-length genes and does not call 25 sequences; 16 of which sit in a *vanG* operon. Among the catalytic domain-only sequences, AMRFinderPlus fails to call 58% of sequences (Fig. 1e). Incorporating the remaining two criteria; i.e presence of an N-terminal transport domain and *vanG* ligase, is integral to resolve both the domain-only architecture and the disagreements between methods (Fig. 1f). First, 276 of the 1,218 domain-only loci carry the missing transport domain as a separate, immediately adjacent gene, and 264 of those also sit beside a *vanG* ligase (Fig. 1f), forming a complete operon split across two genes. This architecture was previously identified in the *vanL* operon of *Enterococcus faecalis*, where *vanT* is split into *vanTm*_*L*_ and *vanTr*_*L*_^10^. The domain-only genes lie no closer to contig ends than full-length ones (6.2% versus 5.8% within 100 bp of an end), thus this result is not due to truncated contigs. Unfortunately, we are unable to confirm 372 of AMRFinderPlus’s 1,111 active-site-positive calls as 227 encode only the catalytic domain within a *vanG* operon without a detectable transport partner, 83 encode the catalytic domain alone with neither a transport partner nor a *vanG* operon (Fig. 1f), and 62 possess a transport function (53 full-length and 9 split domain-only) but sit on contigs too short to evaluate. Crucially, of the 731 active-site-positive loci not called by AMRFinderPlus, 644 lack a transport domain or a *vanG* gene (Fig. 1f). Thus, leaving only 87 genuine misses.

Evaluating all 49,389 CARD predictions confirms that the vast majority of homology calls are orphans lacking both a *vanG* operon or other van genes, independent of gene length (Fig. 2a). When we consider *vanT*_*G*_ in close proximity of a *vanG* ligase, all three approaches converge: 1,491 CARD calls, 985 AMRFinderPlus calls, and 1,085 active-site-positive sequences (Fig. 2a). Conversely, a total of 23,893 CARD calls (8,301 full-length and 15,592 domain-only) sit completely alone with no other van gene in the genome and another 24,005 sit in genomes with other van genes but no *vanG* ligase (Fig. 2a), giving more credence to the idea that almost all uncontextualised CARD calls are misannotated alanine racemases.

**Fig. 2.**
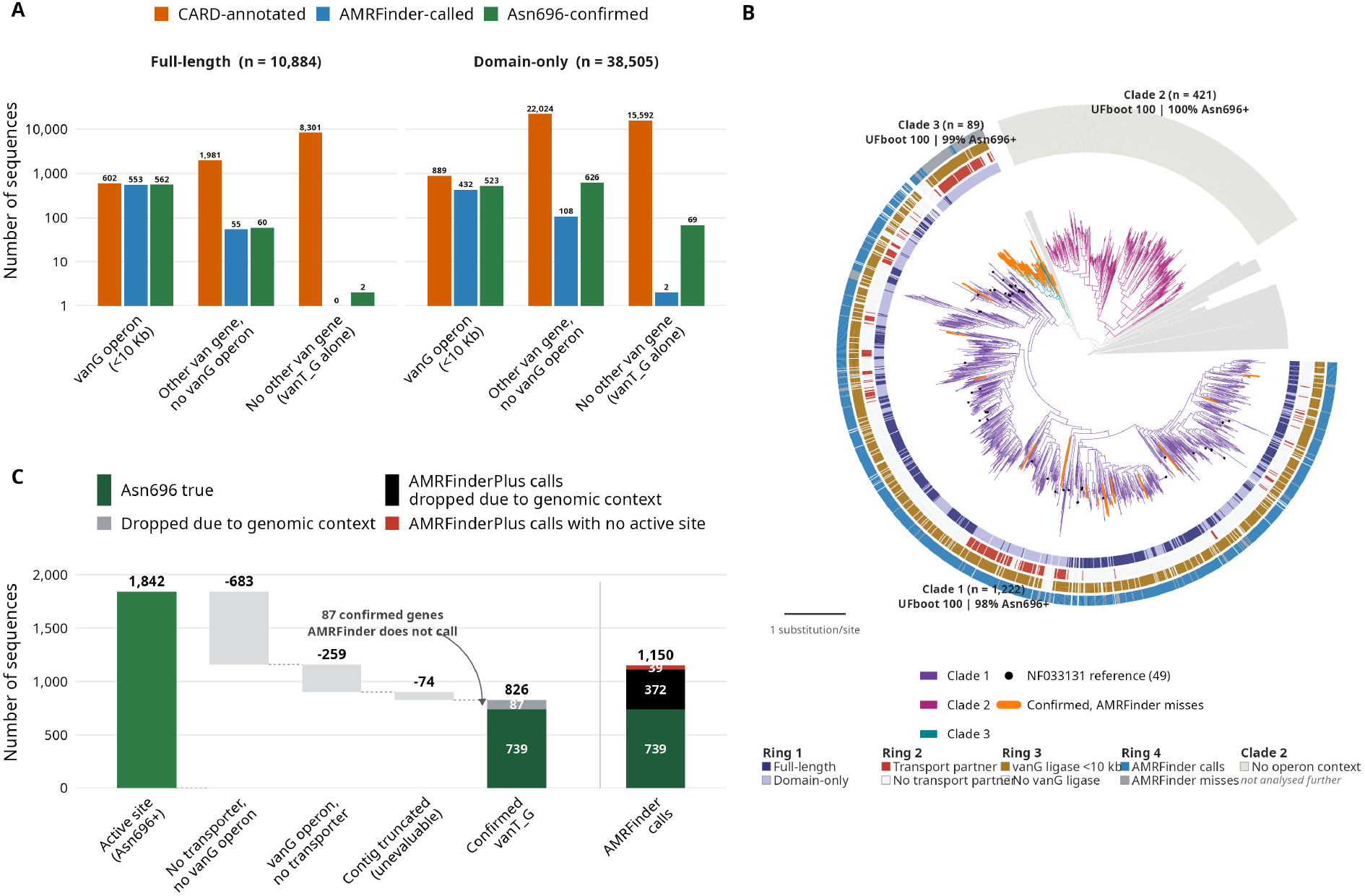
Genomic context and phylogeny separate functional *vanT*_*G*_ from homology artefacts. A) All 49,389 CARD *vanT*_*G*_ calls split by architecture and separated based on the three prediction tools. B) Maximum-likelihood phylogeny of the catalytic domain (7,906 dereplicated sequences; LG+I+R10; midpoint-rooted; scale bar, 1 substitution per site). Branches are coloured by clade, defined as maximal well-supported subtrees (SH-aLRT ≥ 80, UFboot ≥ 95) in which at least 95% of tips carry Asn696. Rings 1–4 annotate Clades 1 and 3: domain architecture; adjacent transport partner; vanG ligase within 10 kb; AMRFinderPlus call. Clade 2 carries no operon context and is shown as a single neutral band rather than four rings. Orange branches mark confirmed genes AMRFinderPlus does not call; black points mark the 49 NF033131 reference proteins, all of which fall in Clade 1. The 6,165 tips in 11 clade-free subtrees are collapsed to grey wedges. C) Reduction from 1,842 active-site-positive sequences to 826 confirmed *vanT*_*G*_, with the reason for each removal, and the composition of the 1,150 AMRFinderPlus calls for comparison.

We placed the 1,842 sequences with Asn696 at the catalytic site along with 6,064 randomly picked CARD called *vanT*_*G*_ sequences on a maximum-likelihood tree (Fig. 2b). Clades were defined as maximal, well-supported subtrees (SH-aLRT ≥ 80, UFboot ≥ 95) with ≥95% Asn696-positive tips to recover three lineages. Clade 1 (1,222 tips) mixes full-length and domain-only sequences, sits close to a *vanG* ligase in 84% of cases, and AMRFinderPlus calls 1,081 sequences. This clade also contains all 49 NF033131 reference proteins. Within Clade 1, AMRFinderPlus calls confirmed full-length and split domain-only genes at nearly identical rates. Clade 2 (421 tips) is almost entirely domain-only, lacks both an adjacent transport domain and a *vanG* ligase in every member, is never called by AMRFinderPlus. Sequences within this clade have substituted arginine or lysine at position 695 adjacent to the active site, where VanTG carries Thr695. Contrastingly, Clade 3 (89 tips) holds the bulk of the *vanT*_*G*_ sequences missed by AMRFinderPlus (Fig. 2b). It is composed exclusively of catalytic domain-only sequences, 80% of which have an adjacent transport partner and 94% a *vanG* ligase, and is taxonomically coherent, with 97.8% of its members belonging to Erysipelotrichales (chiefly *Allobaculum, Faecalibaculum, Longicatena*, and *Ileibacterium*), yet AMRFinderPlus calls only 5 of its 89 sequences. Examining physical distances across the contigs confirms that virtually all operonic *vanG* ligases sit well within the 10 kb window, and reveals a more compact architecture in Clade 3 (Supplementary Fig. 1). The separate transport partner sits at the same distance from *vanTG* in Clade 3 as in other domain-only clades (median 1,118 bp versus 1,087 bp), but the *vanG* ligase sits roughly twice as close to *vanT*_*G*_ in Clade 3 (median 1,119 bp) (Supplementary Fig. 1).

CARD suffers from a substantial overprediction problem that can be immediately resolved using genomic context without requiring new reference data. Applying all three criteria to the 1,842 active-site-positive sequences removes 683 with neither a transporter nor close to a *vanG* ligase, 259 close to a *vanG* ligase, but with no detectable transporter, and 74 on contigs too short to evaluate, leaving 826 confirmed *vanT*_*G*_ genes (Fig. 2c). Of these, AMRFinderPlus recovers 739 but misses 87. These misses are heavily concentrated in a novel split-architecture Erysipelotrichales lineage, which account for 72% of all misses, in Clade 3. Because all 49 NF033131 seed proteins sit in Clade 1 and none in Clade 3, current curated models lack representatives for this gut-associated lineage. This architecture is a genuine and recurrent mechanism of *VanT*_*G*_ function and previously documented only in the *vanL* operon. Adding Erysipelotrichales representatives to reference models will directly improve prediction coverage, pending biochemical validation.

---

**Supplementary Fig. 1:**
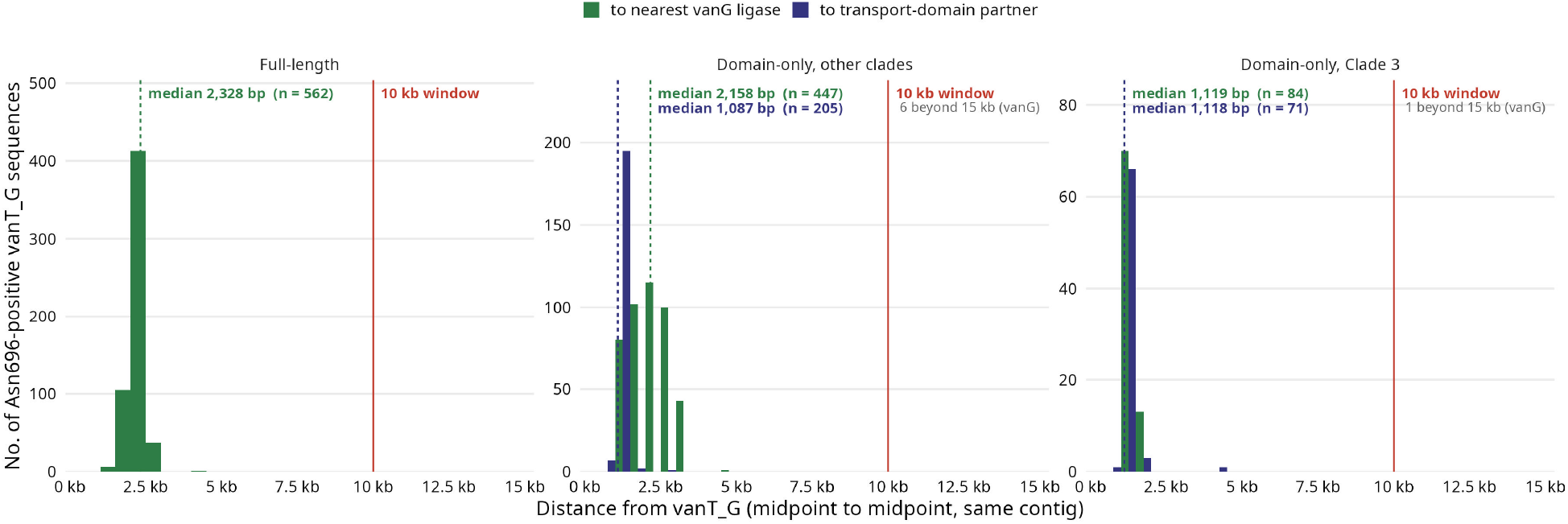
Distance from *vanT*_*G*_ to the genes that define its operon context. Distance from each Asn696-positive *vanT*_*G*_ locus to the nearest *vanG* ligase (green) and, for domain-only loci, to its transport-domain partner (indigo), measured midpoint to midpoint on the same contig and binned at 500 bp.

## Data availability

Workflows and R scripts are available from the first author on reasonable request. All data will be made available upon peer reviewed publication.

## Author contributions

PS, RW and GM conceived the study. RW and PS analyzed the data; PS, RW and GM contributed to interpreting the data and conceptualizing the results. All authors wrote and revised the paper.

